# Tracking ligand-binding-induced structural populations in T4 lysozyme by time-resolved serial crystallography

**DOI:** 10.64898/2026.03.26.714466

**Authors:** Maria Spiliopoulou, David von Stetten, Andreas Prester, Eike C. Schulz

## Abstract

Ligand binding has been shown to induce significant alterations in the conformational landscape of proteins. Traditional crystallography approaches have provided valuable input about the end states in ligand-binding reactions. However, dynamical relationships between ligand binding and backbone rearrangement often remain obscured by crystallographic structures. In the present study, we use time-resolved serial synchrotron crystallography (TR-SSX) to directly visualize indole binding in the cavity of T4 lysozyme L99A in microcrystals under controlled environmental conditions. By integrating fixed target crystallography with LAMA-based ligand delivery, we have been able to track the progression of ligand binding and backbone rearrangement. By utilizing an occupancy refinement protocol, we have been able to quantify structural populations. Our studies reveal that ligand binding for this protein cavity follows a diffusion-limited process that progressively rearranges the *F* -helix of the protein towards a dominant conformational state. These findings establish an observable link between ligand diffusion, occupancy evolution and conformational adaptation within a crystalline environment. More broadly, our work shows how TR-SSX can quantify ligand and conformational populations during binding, providing a framework to interpret structural adaptation in real time.

## 1 Introduction

Bacteriophage T4 lysozyme (T4L) is a small, globular protein consisting of 164 amino acid residues and a molecular weight of approximately 18.6 kDa. It functions as a hydrolase that cleaves the *β*-1,4 linkage between N-acetylmuramic acid and N-acetylglucosamine in the bacterial peptidoglycan, leading to lysis of the bacterial cell wall. Owing to its modest size, well-defined tertiary structure, and extensive mutational characterization, T4 lysozyme has long served as a model system for exploring the relationship between protein structure, dynamics, and function ^1–3^. Beyond its biological role, T4L provides an experimentally accessible framework for studying conformational flexibility in small globular proteins. Numerous experimental and computational studies have shown that intrinsic motions within T4L, particularly in the *F* -helix and surrounding residues, are tightly coupled to its catalytic activity and stability ^2,3^. The size along with the stability of its structure, make the protein an excellent candidate for the application of high-resolution techniques, including X-ray crystallography^4^, nuclear magnetic resonance (NMR) spectroscopy ^5^, and molecular dynamics (MD) simulations ^6^, to gain insight regarding the role of local dynamics and helix motions in the enzymatic activity of the protein.

Among the engineered variants, the L99A mutant has been of particular interest. Replacement of the leucine residue in position 99 with alanine generates an internal, non-polar cavity of approximately 150 Å^3^ that can accommodate benzene and a range of small hydrophobic ligands^2,3,7^. This mutant constitutes a model system for studying protein-ligand interactions. It provides a simple, yet very instructive system for studying the relationships between binding energetics, hydrophobic packing, and conformational flexibility.

Dynamic rearrangements within T4L are not limited to the engineered cavity but also play crucial roles in its native catalytic function. Motions of the *F* -helix and neighboring regions facilitate the threading and translocation of peptidoglycan fragments through the active-site cleft, highlighting the importance of backbone flexibility for substrate processing and enzymatic turnover^8,9^. Collectively, these studies have established T4L as a benchmark system to investigate how local and global conformational transitions enable both catalysis and ligand binding.

Recent room temperature (RT) structural studies of T4L-L99A provide further support for the dynamic nature of its binding cavity; for example, the RT X-ray structure of L99A soaked with toluene (PDB ID: 7L39) reveals backbone and side chain conformations not observed under cryogenic conditions ^10^. These observations support the idea that critical functional states can be obscured or entirely suppressed within cryogenic frameworks.

Understanding the structural basis of ligand binding has been a fundamental objective in the field of protein science, as it is directly linked to molecular recognition, catalysis, and allosteric regulation. Conventional crystallography has provided atomic level snapshots of ligand bound and unbound states, yet the dynamic sequence of conformational changes that connect these states often remains inaccessible^11^. Time-resolved X-ray crystallography (TRX) has emerged as a powerful method to bridge this gap by directly visualizing structural intermediates and reaction processes at temperatures close to physiological conditions^12–14^.

In the past decade, this field has undergone a resurgence, primarily sparked by the introduction of X-ray free-electron laser (XFEL) sources and the development of serial crystallography techniques^15–17^. In particular, serial crystallography has become a central implementation of TRX, enabling data collection from large numbers of microcrystals while minimizing radiation damage and allowing measurements at room temperature^18–20^. The successful application of serial crystallography at XFELs was quickly adopted at synchrotron facilities, which can provide equivalent data quality for sufficiently large crystals^20–23^.

While serial synchrotron crystallography (SSX) at monochromatic MX beamlines cannot reach the ultra-fast timescales that are accessible to XFELs, they are, however, the obvious choice when it comes to studying enzymatic mechanisms beyond the microsecond time domain^24^. Their comparably wide distribution enables studies of a larger number of model systems and supports the development of new techniques that will benefit the field.

The recent developments in time-resolved serial synchrotron crystallography (TR-SSX) have been analyzed in several reviews ^16,25–31^. On a methodological level, time-resolved room temperature crystallography is capable to capture protein dynamics on biologically relevant timescales. For instance, slow relaxation processes on the order of seconds have been visualized by time-resolved oscillation crystallography at synchrotron sources ^32^. Such methods enable studies of ligand binding that go beyond static structures. Through rapid reaction initiation, typically via laser excitation, TRX enables capture of transient, out-of-equilibrium conformations without the need for engineered mutant traps. Reaction initiation can also be achieved by ligand delivery using microdroplet injection systems: the LAMA (*liquid application method for time-resolved analyses*) technique allows *in situ* deposition of ligand solution onto microcrystals within a serial synchrotron crystallography (SSX) framework^13^, while the HARE (*hit-and-return*) method enables versatile delay times (ms - minutes) by revisiting the same crystal position after controlled delay times^12^. The combination of serial data acquisition with LAMA-based ligand delivery and HARE timing provides a versatile platform for monitoring ligand binding and associated conformational changes in real time, avoiding artifacts from cryogenic trapping or mutation-based stabilization.

T4 lysozyme L99A (T4L-L99A) has served as a widely studied model for dissecting ligand binding within a buried hydrophobic environment. Early crystallographic analyses revealed that ligand association involves subtle but functionally important rearrangements of the loops surrounding the cavity. In particular, a halogen bond formed by Met102 contributes to stabilizing halogenated ligands and modulating loop conformations^33^. These findings underscore how even minor side-chain substitutions in the hydrophobic core can influence both ligand accommodation and the conformational landscape of the protein.

Subsequent studies demonstrated that the volume and shape of the L99A cavity varies systematically with ligand size. Crystallographic analysis of benzene and its derivatives revealed a clear correlation between ligand dimensions and cavity expansion^34^. However, these investigations were conducted using cryocooled crystals, capturing static end-states rather than the continuous range of conformations accessible under physiological conditions.

More recent work has emphasized the inherent plasticity of the L99A cavity, revealing that ligand binding induces both local and global adjustments within the protein scaffold ^35^. This concept challenges the notion of the cavity as a rigid, preformed site, instead suggesting that the protein backbone actively participates in binding-induced conformational changes. Despite these advances, the temporal structural details of these rearrangements remain poorly characterized. In the present study, we seek to provide further insight into this process by time-resolved crystallography, to elucidate the structural and dynamical basis of ligand-induced conformational changes in T4L-L99A.

## 2 Results and Discussion

### 2.1 Indole binding expands the hydrophobic cavity

To evaluate ligand-induced conformational changes in T4L-L99A, crystals were statically soaked with indole to a final concentration of 0.05 M and measured by serial data collection at 20°C. The resulting structures confirm indole binding within the engineered hydrophobic cavity (**Fig. 1**). Comparison with the apo dataset reveals distinct backbone rearrangements, most prominently a displacement of approximately 1.7 Å in the *F* -helix region adjacent to the binding site (**Fig. 1**). Closer inspection of the cavity environment shows that residues lining the surrounding helices exhibit positional shifts that correlate with the degree of cavity opening, suggesting an adjustment of the local backbone to accommodate ligand entry and binding. This rearrangement highlights the intrinsic plasticity of the cavity region and its role in facilitating ligand-induced conformational adaptation resulting in an open *F* -helix state.

**Fig. 1.**
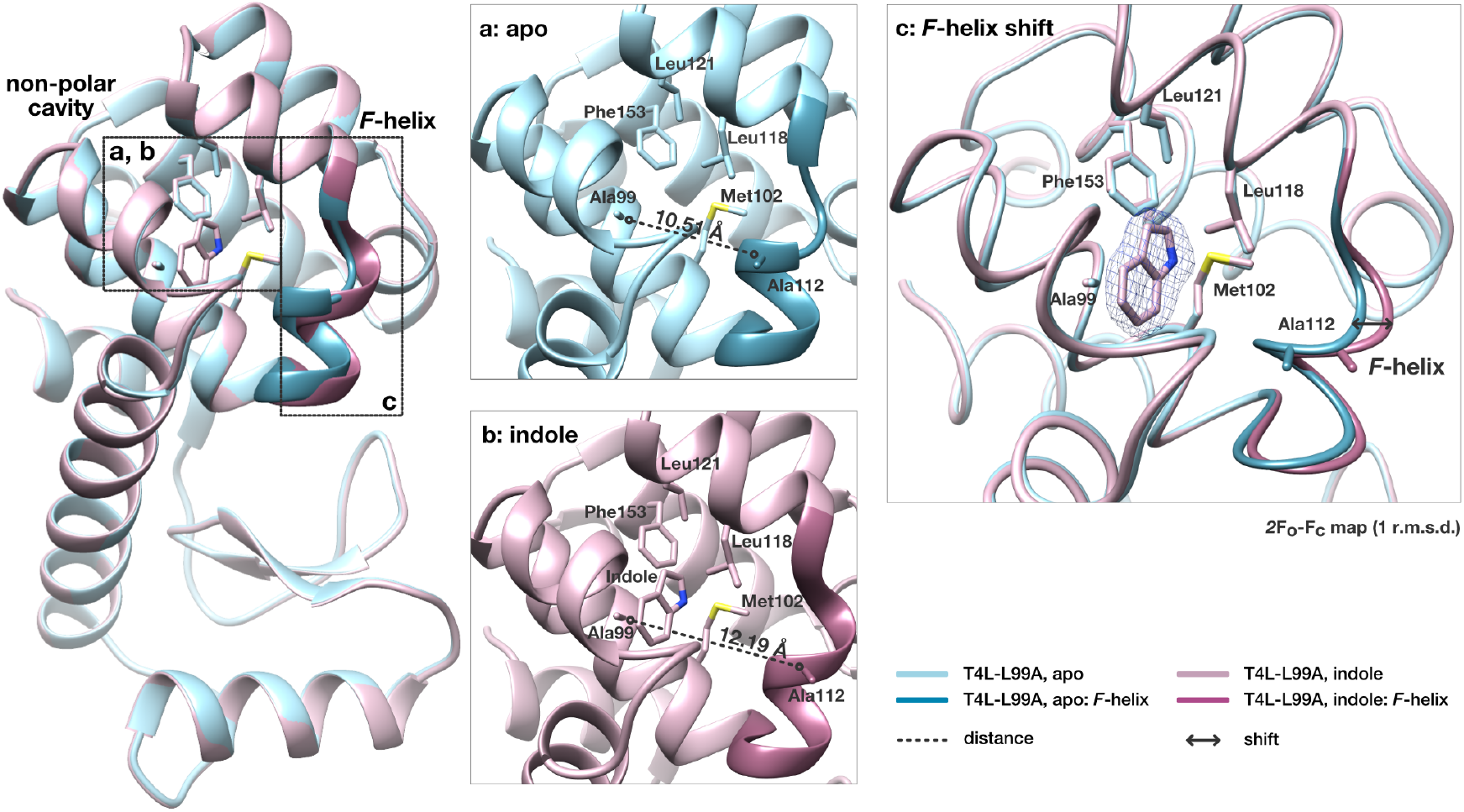
Indole-induced backbone changes in T4L-L99A. Ribbon representation of the non-polar cavity **(a, b)** in the absence (blue) and presence of indole (pink) after static soaking. The conformational change of the *F* - helix **(c)** can be observed for the bound state of T4L, leading to a further opening of the cavity of about 1.7 Å, based on the Ala99 to Ala112 distance.

### 2.2 Ligand binding modulates temperature-dependent flexibility

Serial crystallographic datasets for both the apo and indole-bound forms of T4L-L99A were collected across a temperature series (10–30°C) at comparable resolution, enabling a consistent evaluation of temperature-dependent structural changes. Global C*α* RMSD analysis ^36,37^revealed a clear distinction between the two states: the apo protein exhibited progressively larger deviations with increasing temperature, whereas the indole-bound form remained comparatively stable (**Fig. 2 a, b**). This trend was further reflected in the atomic displacement parameters, which increased substantially in the apo structures and particularly within loop regions surrounding the engineered cavity, while the ligand-bound protein showed a markedly reduced temperature sensitivity (**Fig. 2 c, d**). These observations indicate that indole binding exerts a stabilizing influence on the local backbone, limiting the thermally driven structural fluctuations that are prominent in the ligand-free state.

**Fig. 2.**
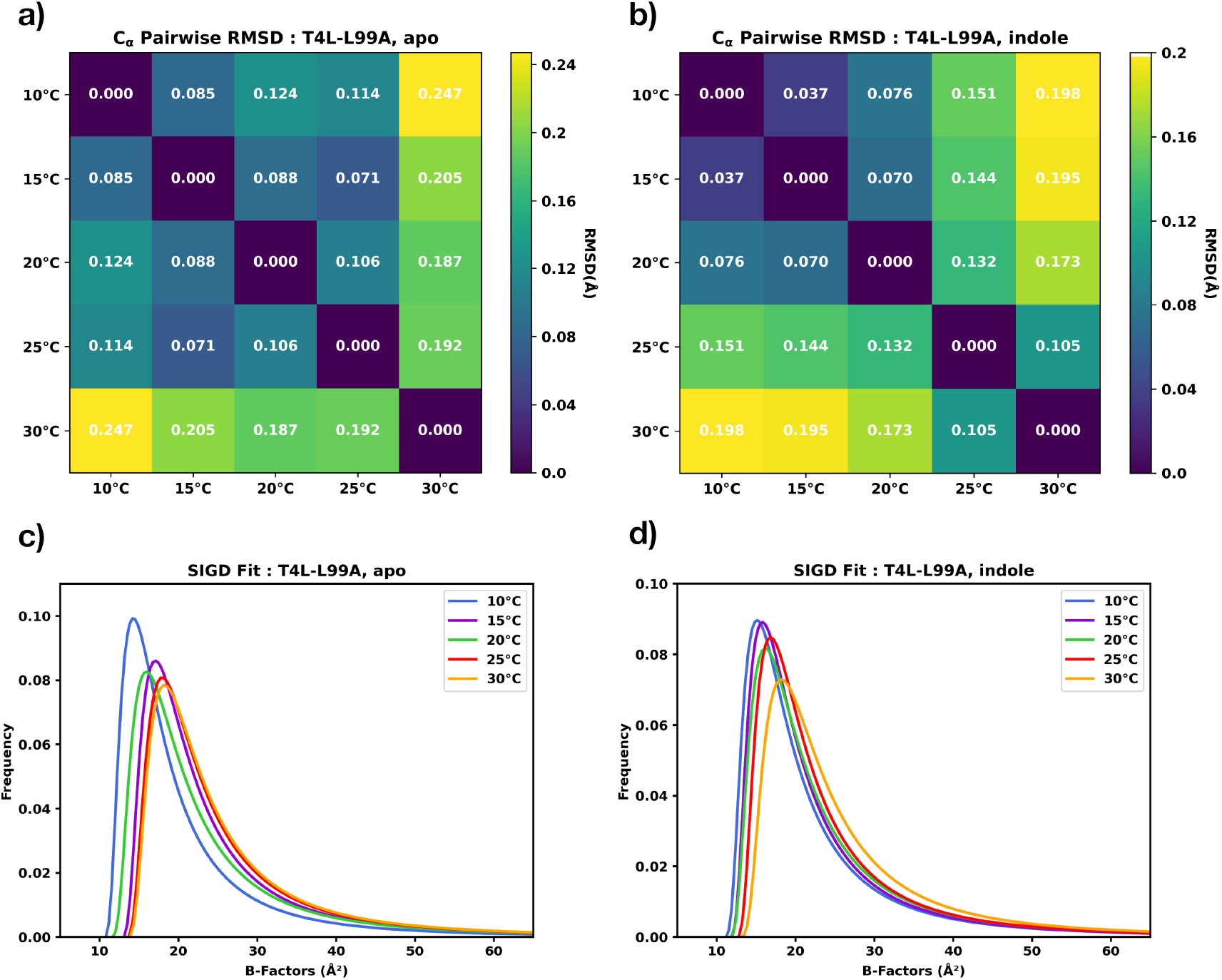
Temperature-dependent structural deviations for T4L-L99A. C*α*-RMSD analysis **(a, b)** and atomic displacement parameters fitted to a shifted inverse gamma distribution **(c, d)** for apo and indole-bound protein.

To further assess conformational variability, a torsion-angle–based representation of protein entities (RoPE) analysis^38^ was performed for the apo and indole-bound datasets across the full temperature series (**Fig. 3**). In the absence of ligand, RoPE projections showed a progressively wider dispersion with increasing temperature, indicating that torsion-angle heterogeneity emerges across multiple regions surrounding the engineered cavity. By contrast, the indole-bound structures remained more compact in RoPE space, displaying a focused shift dominated by torsion-angle deviations within the *F* -helix adjacent to the binding cavity. This distinction suggests that ligand binding constrains the conformational landscape, stabilizing much of the protein while selectively modulating the flexibility of the *F* -helix. Taken together, the RoPE analysis highlights differences in the dynamic response: the apo protein samples a broader set of conformational states at increasing temperatures, whereas indole binding restricts this expansion and channels structural adaptation primarily into the *F* -helix.

**Fig. 3.**
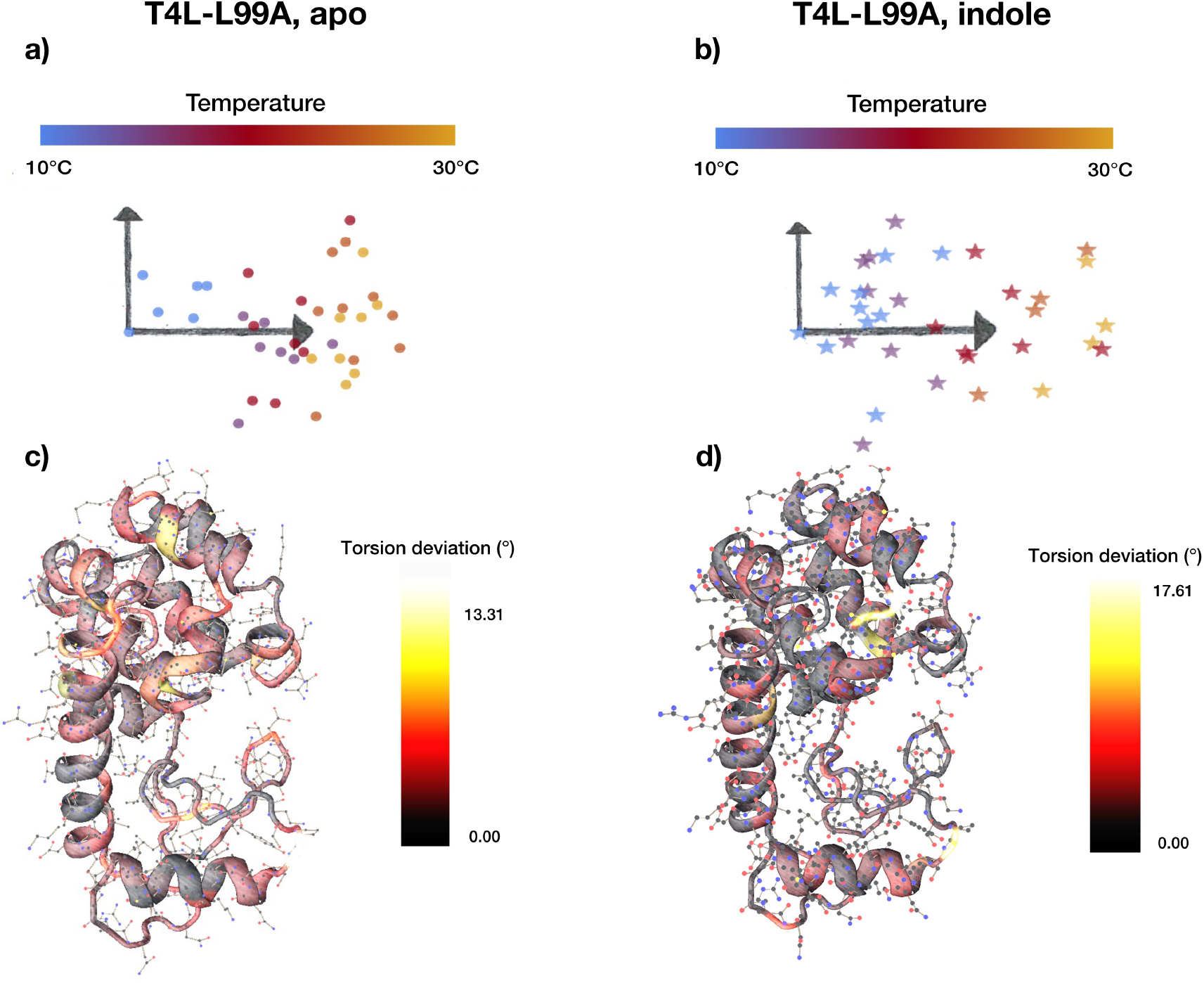
RoPE analysis. **(a, b)** RoPE plots based on the torsion-angle deviation for both protein states across temperatures. Each point represents a structure projected onto the first two principal components. **(c, d)** Corresponding torsion-angle deviations mapped as backbone heatmaps, highlighting local structural responses to temperature changes.

### 2.3 Quantitative analysis of indole binding

#### 2.3.1 Occupancy determination and conformational sampling

In order to quantify the ligand binding and the dynamics of the *F* -helix, T4L-L99A microcrystals of size 15×15×15 *µ*m were mounted on fixed-target SSX chips and exposed to a 0.05 M solution of indole using the HARE & LAMA methods^13,14^, allowing precise time-resolved sampling at well-defined time delays from 0.5 s up to 40 s. The measurements were performed in a controlled environment of 20°C and 95 % relative humidity within the environmental control box of the T-REXX endstation^39^ (**Fig. 4**).

**Fig. 4.**
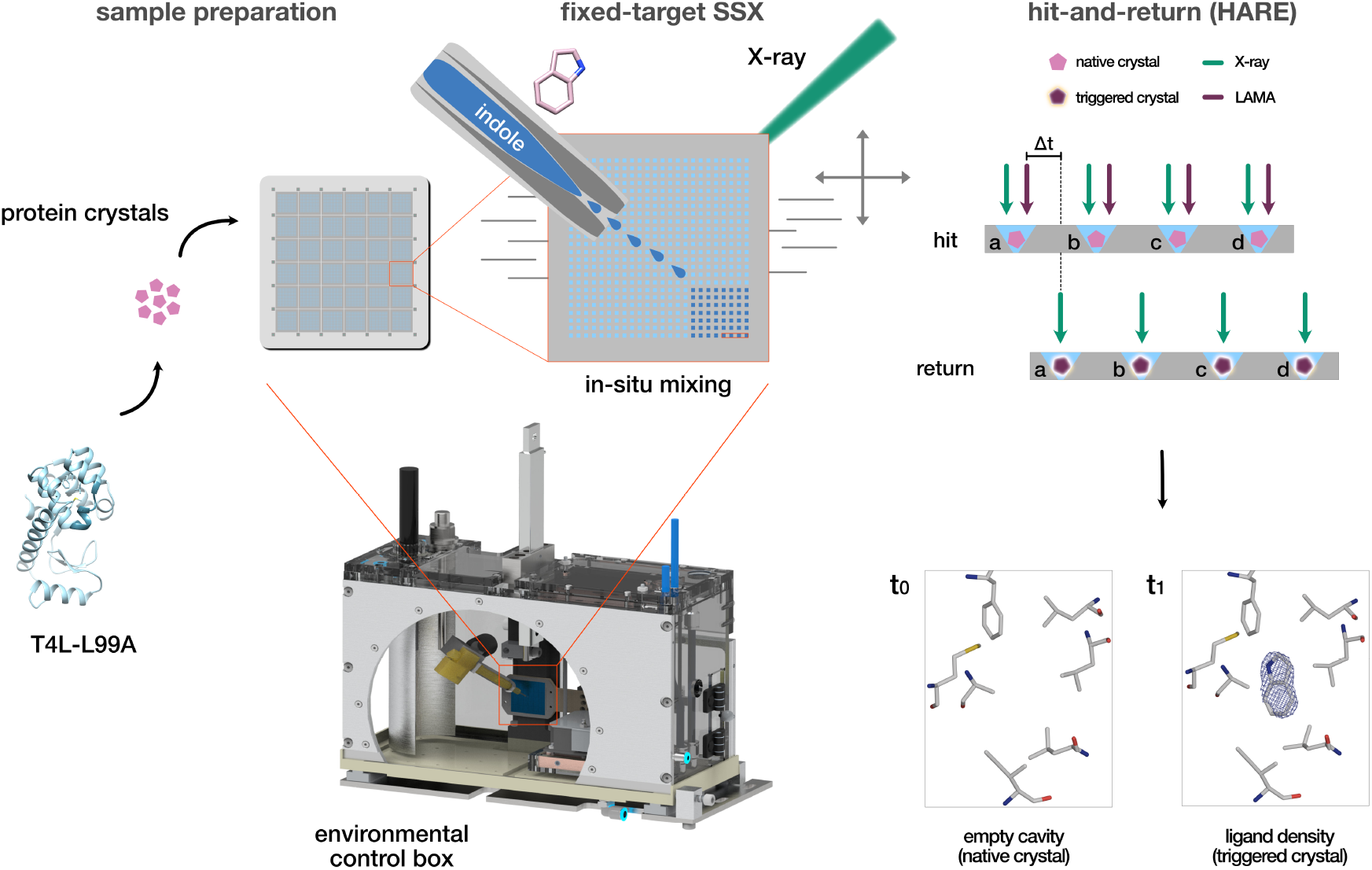
Experimental process for the time-resolved T4L-L99A–indole binding.

The 8 obtained protein structures were refined until convergence using `phenix.refine` and manual refinement in coot **(Supplementary Table 1)**. For a more accurate determination of the occupancies and *B* -factors, these results were analyzed using a batch refinement method. For each individual protein structure, 500 independent refinement runs were performed with randomized initial occupancies and *B* -factors of indole and the two (open and closed) *F* -helix conformations. This allows the quantification of the refined occupancies, which include the inherent uncertainties in the calculation of the occupancies of the diffusion-limited binding events. The scatter plots in **Fig. 5** show clear clusters of the indole and the two *F* -helix states, where the helix predominantly adopts one of them for low or high indole occupancies. The refined occupancies are summarized in **Supplementary Table 1** and the occupancy plots for all time points are illustrated in **Supplementary Figure 1**.

**Fig. 5.**
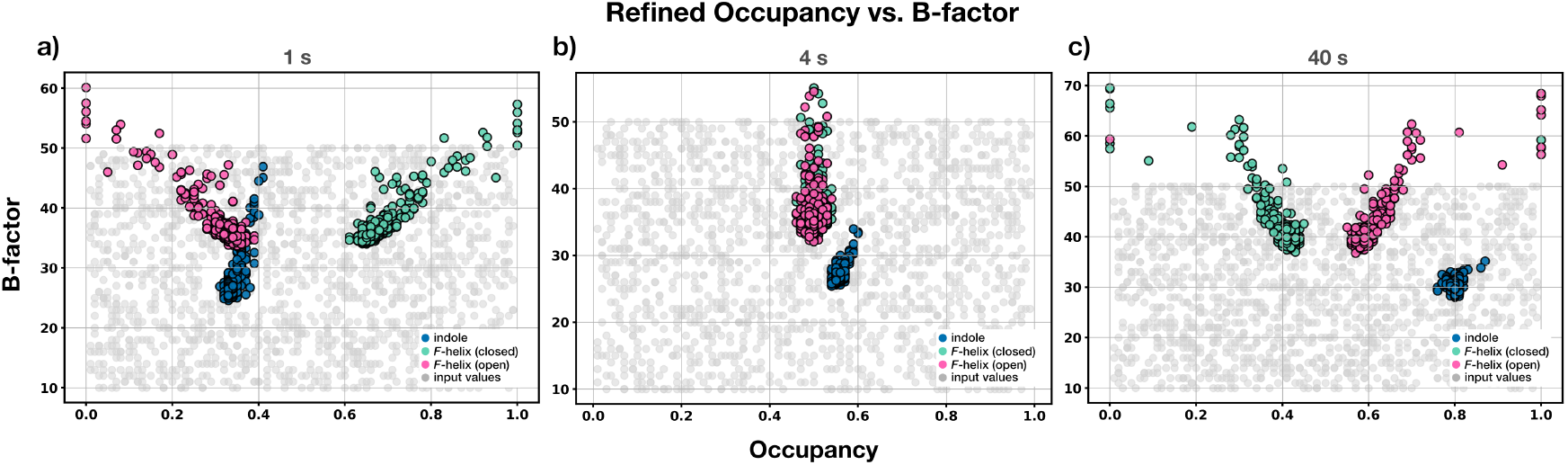
Occupancy–*B* -factor distributions from independent refinements. Scatter plots of refined occupancies versus *B* -factors for indole (blue) and the *F* -helix in open (green) and closed (pink) conformations for certain time points: 1 s **(a)**, 4 s **(b)**, and 40 s **(c)**. Each point represents one of 500 independent refinement runs with randomized starting parameters; grey points indicate initial randomized values prior to refinement. Distinct clusters highlight the refinement behavior and occupancy convergence of each group in consistence with increasing indole and open *F* -helix conformations, the occupancy of the closed *F* -helix conformation decreases over time.

#### 2.3.2 Crystalline population sampling

Inspection of the unit cell parameters refined with `CrystFEL` revealed a systematic change along the *c* axis during ligand binding. While the apo room-temperature structure exhibited a *c* axis length of 97.1 Å, datasets collected at longer delay times, corresponding to higher refined indole occupancies, showed an increase towards approximately 97.9 Å (≈ 0.8%). Changes in the unit cell dimensions have already been observed for T4L-L99A in complex with indole after 1 and 10 s delay time for cryogenic conditions^40^.

Our results demonstrate that the *c* axis shift was not abrupt but appeared progressive across the time series. Notably, for intermediate time points, the distribution of *c* axis values displayed two distinct populations: one centered near 97.1 Å and a second near 97.9 Å. The relative contributions of these populations changed as a function of time (**Supplementary Figure 2**). Early delay points were dominated by the shorter unit cell parameter, whereas later time points showed an enrichment of the expanded lattice. Intermediate delays exhibited comparable contributions from both populations, suggesting coexistence of two crystalline states during the binding process.

To further investigate this behavior, we sorted the datasets based on the refined *c* axis parameter. Diffraction patterns with c ≤ 97.5 Å were grouped into a ‘small-cell’ population, whereas patterns with c ≥ 97.5 Å were assigned to a ‘large-cell’ population in two distinct processing pipelines. The variation of the unit cell dimensions from the initial value (*a*_0_=*b*_0_=60.9 Å, *c*_0_=97.1 Å; total, small-cell, *c*_0_=97.9 Å; large-cell) is described in **Fig. 6 a-c**. This separation enabled a population-specific structural analysis to assess how lattice expansion correlates with ligand occupancy and conformational redistribution.

When comparing the number of indexed hits obtained from the two processing strategies across time, opposite trends were observed: the fraction of hits assigned to the large-cell population increased with time, whereas the fraction corresponding to the small-cell population decreased (**Fig. 6 d**). The temporal evolution of these fractions could be approximated by a four-parameter logistic (4PL) fit. Notably, this trend follows the same qualitative behavior observed for the refined occupancies of indole and the two *F* -helix conformations, with increasing populations associated with indole binding and the open *F* -helix state, and a corresponding decrease in the closed state, as described below (**Fig. 7**).

**Fig. 6.**
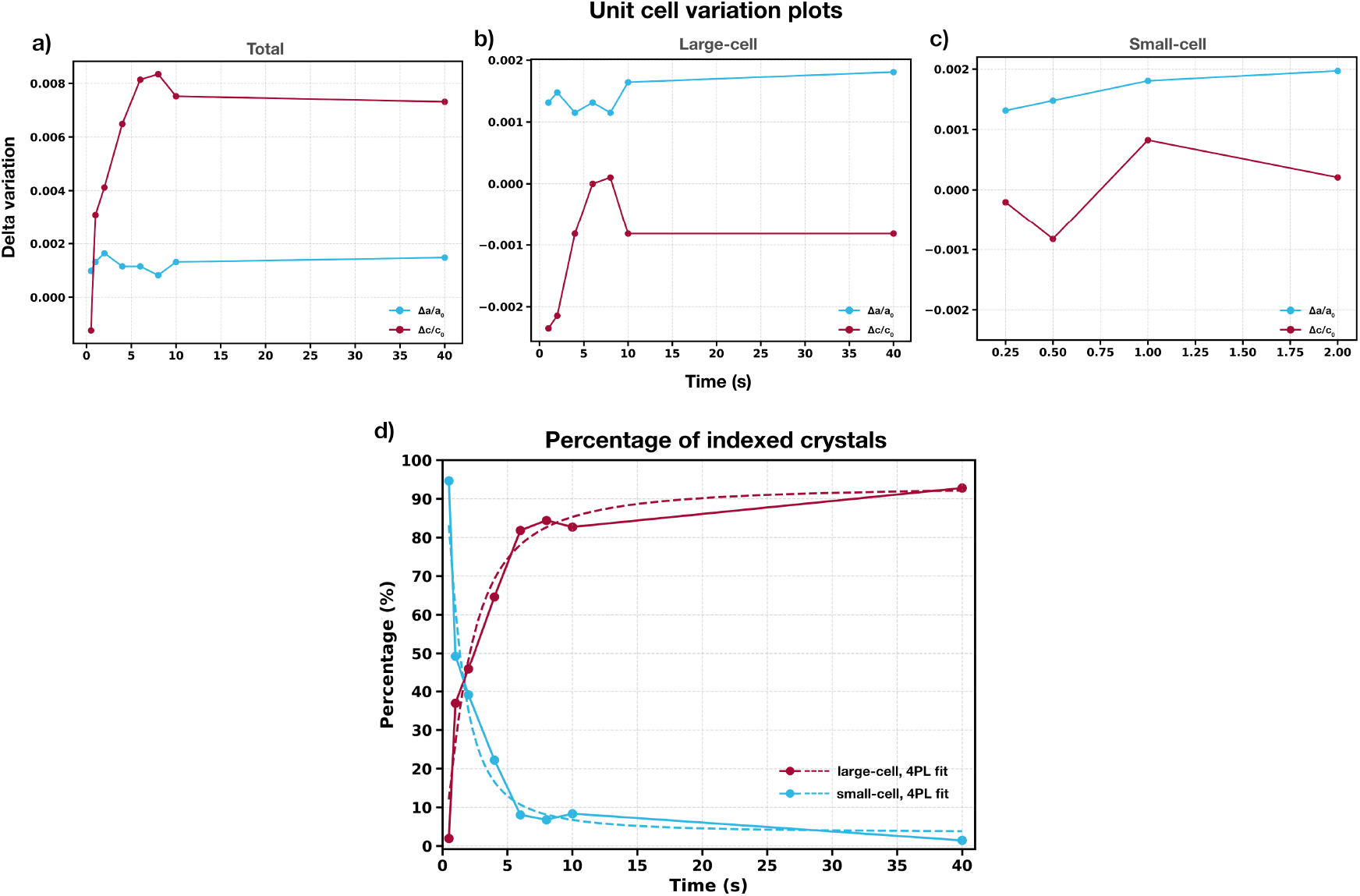
Unit cell variations and percentage of indexed crystals. Scatter plots illustrating the delta variations in unit cell parameters Δ*a*/*a*_0_ (blue) and Δ*c*/*c*_0_ (violet red) of the T4L-L99A refined cell as a function of time for the total **(a)**, large-cell **(b)** and small-cell **(c)** processing. **(d)** Percentage of the indexed crystals for the large-cell (violet red) and small-cell (blue) processing and the corresponding 4PL fit in dashed lines (*R*^2^=0.962; large-cell, *R*^2^=0.950; small-cell).

**Fig. 7.**
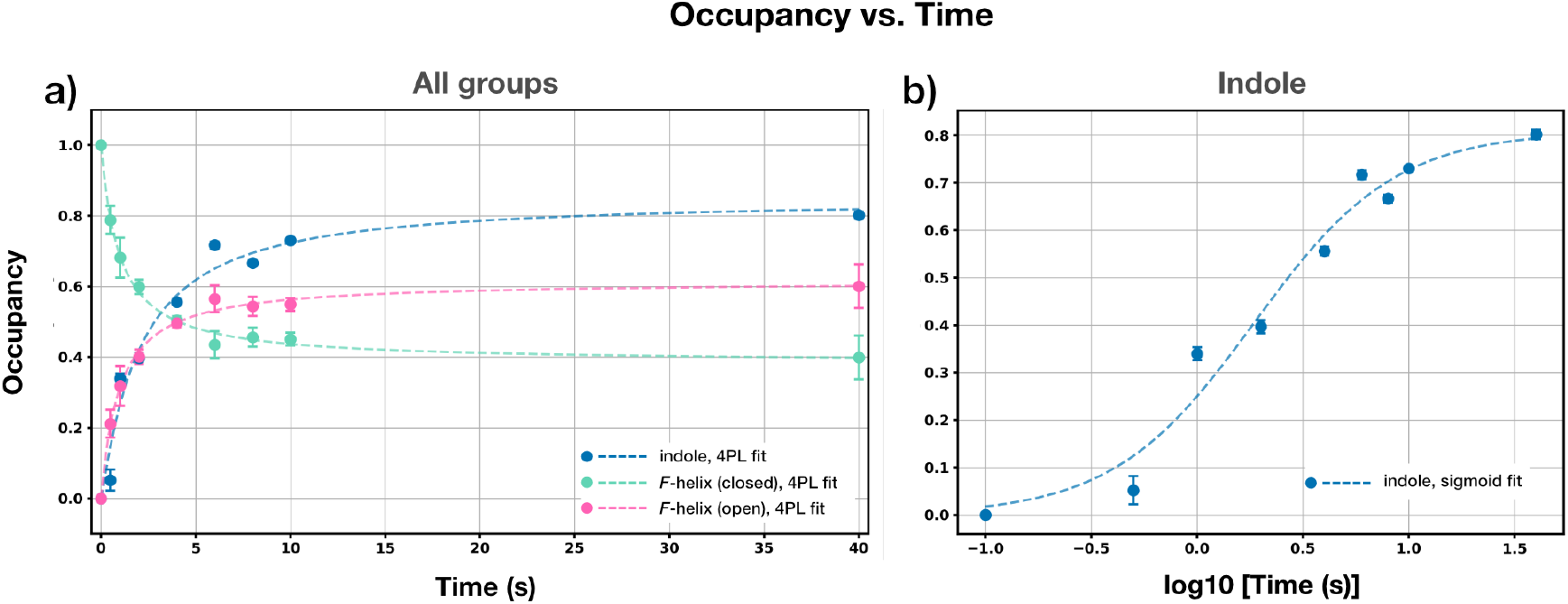
Occupancy over time plots. Occupancy values extracted as a function of time for indole (blue) and *F* -helix in open (green) and closed (pink) conformations based on occupancy–*B* -factor analysis. Data points represent fitted occupancies, with error bars indicating standard deviations, and dashed lines corresponding to **(a)** four-parameter logistic fit for all groups (*R*^2^=0.988; helix, *R*^2^=0.972; indole) and **(b)** sigmoidal fit for indole (*R*^2^=0.972). The right plot displays the indole occupancy over time in a log scale.

#### 2.3.3 TR-SSX resolves indole binding as function of time

TR-SSX allowed direct visualization of indole binding to T4L-L99A under diffusion-limited conditions. Using microcrystals of approximately 15×15×15 *µ*m, ligand binding into the non-polar cavity was visualized through the gradual increase in the refined occupancy of indole. At early delay times, partial occupancies were observed, consistent with ligand diffusion through the solvent channels of the crystal lattice and initial engagement of the cavity, as predicted for diffusion-controlled reactions in microcrystalline samples^41,42^.

With increasing delay, occupancies approach saturation, indicating completion of the binding process within a timescale mainly determined by the crystal environment and the applied indole concentration (0.05 M), rather than by intrinsic chemical steps, consistent with prior TR-SSX studies highlighting the impact of microcrystal solvent channels and ligand delivery conditions on binding^27,43,44^. Notably, the progressive increase in indole occupancy is correlated by a shift in the *F* -helix towards predominantly one of the two modeled conformations (open or closed), indicating the structural adaptation of the protein (**Fig. 5**). The refined occupancies for indole and the two *F* -helix conformations (open/close) are plotted as a function of time (**Fig. 7**), illustrating the progressive binding of the ligand and the accompanying stabilization of the helix into its preferred conformation. This can also be observed in the increasing density of the ligand in the cavity (**Fig. 8**).

**Fig. 8.**
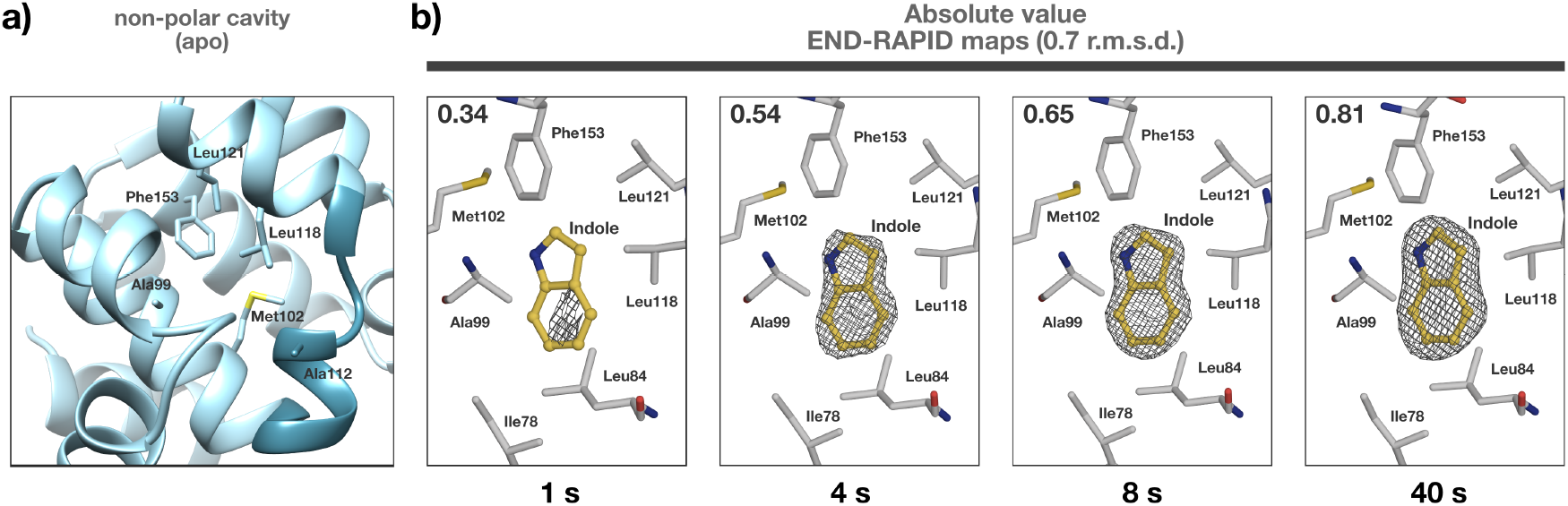
Indole binding in the non-polar cavity of T4L-L99A. **(a)** Closeup of the cavity and the surrounding residues. **(b)** Absolute value END-RAPID maps for indole binding at certain delay times obtained by the HARE approach; the refined occupancies for indole are indicated on the top left for each time point.

Together, these observations establish TR-SSX as a powerful approach for resolving ligand binding in proteins, linking diffusion, occupancy evolution, and functionally relevant conformational changes that are inaccessible to static crystallographic experiments^41–44^.

## 3 Conclusions

In this study, we show the utility of TR-SSX to quantitatively elucidate ligand binding and conformational adaptation in T4L-L99A. By combining fixed-target serial data collection with controlled ligand supply, we monitor the diffusion-limited process of indole ligand binding and relate the increasing ligand occupancy to the adaptation of *F* -helix conformations to a preferred state.

This work expands beyond traditional static crystallographic results, establishing a relationship between ligand diffusion, increasing ligand occupancy, and protein backbone flexibility. Complementary analysis of unit cell parameters further revealed the presence of two crystalline populations distinguished by differences in the c axis. Selective processing of these small- and large-cell states showed systematic changes in their relative population over time, following a trend comparable to the observed ligand occupancy and *F* -helix conformational sampling. These observations clearly link crystallography with ligand binding.

The combination of TR-SSX data collection methods with quantitative refinement approaches thus provides a general and versatile tool to address transient ligand-binding phenomena and conformational adaptation in proteins, opening new possibilities to investigate dynamic structural processes.

## 4 Materials and methods

### 4.1 Protein purification and crystallization

T4L-L99A mutant lysozyme gene was cloned in pET29b(+) vector (Genscript) and expressed in BL21 DE3 *E. coli* cells grown in LB medium supplemented with 50 *µ*g/mL kanamycin at 37°C until an OD600 of 0.7–0.8 was reached. Protein expression was induced by addition of IPTG to a final concentration of 0.001 M and the cells were further incubated at 16°C for 18 hours. The cells were harvested by centrifugation (7500×g, 20 min, 4°C) and pellets were stored at 20°C until purification. The pellets were resuspended in lysis buffer (0.05 M NaH_2_PO_4_/Na_2_HPO_4_ pH 6.5, 0.002 M EDTA) and sonicated for lysis.

The protein was purified by ion exchange against 0.05 M NaH_2_PO_4_/Na_2_HPO_4_ pH 5.5, 1 M NaCl, 0.002 M EDTA and size exclusion chromatography with a final buffer of 0.05 M NaH_2_PO_4_/Na_2_HPO_4_ pH 5.5, 0.1 M NaCl, 0.002 M EDTA ^45^.

Batch crystallization was performed in a total volume of 20 *µ*L by mixing equal volumes of protein at 22 mg mL^*−*1^ and crystallization buffer consisting of 4.0 M NaH_2_PO_4_/K_2_HPO_4_ pH 7.0, 0.1 M *1,6* -hexanediol and 0.15 M NaCl (PDB ID: 3K2R).

### 4.2 Calculation of C*α*-RMSD

In order to quantify and visualise the changing global structural differences, a pairwise backbone Root Mean Square Deviation (C*α* RMSD) ^36,37^ was calculated T4L-L99A and represented as a categorical heatmap. Before calculating the RMSD for a pair, the two structures were aligned via Singular Value Decomposition (SVD) ^46^ using the Biopython package `Bio.SVDSuperimposer` in order to minimise the resulting RMSD value ^39^. A temperature dependent clustering is observed for both native and ligand-bound protein.

### 4.3 Beamline experimental setup and X-ray data collection

Diffraction datasets were collected at the P14.2 (T-REXX) beamline of PETRA III (DESY, Hamburg) at the EMBL Hamburg endstation. Data were acquired using an X-ray beam of 10×7 *µ*m on an Eiger 4M detector at 12.7 keV. The data collection procedure followed previously described protocols ^12^ and was performed within the environmental control box installed at the T-REXX endstation^39^. In brief, microcrystals mounted on a HARE-chip solid target containing 20,736 wells were translated through the X-ray beam using a three-axis piezoelectric translation stage (SmarAct, Oldenburg, Germany)^14^. Time delays were generated using the HARE method, while reaction initiation was achieved via in situ droplet injection employing the LAMA technique^12–14^. The HARE approach minimizes overall wall-clock time for extended delay ranges, whereas the LAMA method enables reaction initiation through localized droplet deposition. As a general guideline, approximately 5,000 still diffraction patterns were recorded per time point, as previously determined to be sufficient^23^. Data collection parameters for each individual time point are summarized in **Supplementary Table 1**.

### 4.4 Reaction initiation via the LAMA method

Time-resolved ligand delivery was performed using the Liquid Application Method for time-resolved Analyses (LAMA) ^13^ in combination with a fixed-target serial synchrotron crystallography (SSX) setup. Protein microcrystals were loaded into the wells of the HARE chip, leaving only a thin layer of mother liquor. Freshly prepared, sterile-filtered, and degassed ligand solution was applied via a piezo-driven nozzle (autodrop pipette AD-KH-501; microdrop Technologies GmbH, Norderstedt, Germany) positioned above the chip. Single droplets (75–150 pL) were deposited onto each crystal, resulting in a total ligand consumption of approximately 1.5 *µ*L per chip. Each reaction cycle involved recording a reference image (t_0_), triggering ligand deposition, and acquiring a time-resolved image (t_1_) after the desired delay. Droplet transit time (≈ 0.5 ms) was included in the timing to ensure accurate temporal control. The experimental process is illustrated in **Fig. 4**. This approach enables precise, low-consumption ligand application directly on chip-mounted crystals, compatible with fixed-target SSX measurements^12,14^.

### 4.5 Data processing and structure refinement

Diffraction data were processed with `CrystFEL` package ^47^. Molecular replacement was conducted in `Phaser` using PDB-ID 4W51 as starting model for T4L-L99A ^48^. Structures were refined using iterative cycles of `REFMAC` or `phenix.refine` and `coot`^49–51^. Molecular images were generated in `PyMol` ^52^.

END-RAPID maps were generated in order to examine lower occupancy features maps as they estimate local noise directly from experimental and model-derived errors ^53^. These maps are presented in an absolute scale, and a noise threshold has been redefined, which helps in detecting weak, but real, electron density that may be obscured in conventional maps and enables direct comparison of electron density features, since maps are on an absolute scale.

#### 4.5.1 Sub-dataset splitting and analysis with RoPE

For the RoPE analysis, each dataset was split into subsets of at least 2000 diffraction patterns each using partialator^47^, which were then independently refined with `DIMPLE` ^54^, without human intervention. This allows us to assess the relative contributions of random fluctuation and genuinely temperature-dependent changes within the crystal structure. The output of these PDB files was analysed in RoPE^38^ the top three coordinates were rotated for clarity and projected onto 2D of the atomic coordinate differences, and corresponding temperature metadata.

#### 4.5.2 Unit cell sorting in CrystFEL

For population specific processing, certain steps in `CrystFEL` were modified ^47^. Specifically, separate . `cell` files were generated by setting *c* axis to 97.1 Å (small-cell) and 97.9 Å (large-cell), corresponding to the two observed lattice populations. During indexing, the unit cell tolerance parameter was reduced from the default for cell axes of 5% to 0.41% only for *c* axis. This constraint ensured that only crystals with refined *c* axis values within ± 0.41% of 97.1 Å were included in the small-cell dataset, and correspondingly, only crystals within ± 0.41% of 97.9 Å were included in the large-cell dataset. All other processing parameters remained unchanged. Merging and subsequent structure refinement were performed independently for each population using identical refinement protocols.

#### 4.5.3 Occupancy and ADP refinement

In order to test the robustness of refined occupancies and atomic displacement parameters, a batch-refinement strategy was implemented using a single reference PDB model as input. From this model, 500 derivative structures were generated by randomly assigning initial occupancies (0.01–0.99) and *B* -factors (10–50 Å^2^) to selected groups. The refined groups consisted of (i) the bound indole molecule and (ii) the *F* -helix modeled in open and closed conformations, with their occupancies constrained to a total occupancy of 1.0 during refinement. All other model parameters were kept unchanged. Each derivative model was refined independently using `phenix.refine` with six automated refinement cycles and no manual intervention. The output, being consisting of 500 refined models, was analyzed to extract final occupancies and *B* -factors of the targeted groups, enabling statistical evaluation of refinement behavior under varying initial conditions.

## Acknowledgments

ES acknowledges support by the Federal Ministry of Education and Research, Germany, under grant number 01KI2114. Funded by the European Union (ERC, DynaPLIX, SyG-2022 101071843). Views and opinions expressed are however those of the author(s) only and do not necessarily reflect those of the European Union or the European Research Council Executive Agency (ERCEA). Neither the European Union nor the granting authority can be held responsible for them.

## Author Contributions

M.S. performed protein expression, purification, prepared the protein crystals, performed data collection, analyzed the data and wrote the manuscript; D.v.S. supported the performed measurements, processed and analysed the diffraction data; A.P. established the occupancy batch refinement pipeline; E.C.S. supervised and wrote the manuscript; all authors discussed and corrected the manuscript.

## Data and materials availability

The data that support this study are available from the corresponding authors upon request. All crystallographic data have been deposited in the Protein Data Bank (PDB) under following accession numbers: 29NF, 29NG, 29NH, 29NI, 29NJ, 29NK, 29NL, 29NM, 29NN, 29NO. Further details are available in **Supplementary Table 1**.

## Declarations

There are no competing interests to declare.

