## Supplementary Information for "Tracking ligand-binding-induced structural populations in T4 lysozyme by time-resolved serial crystallography"

#### Supplementary figures

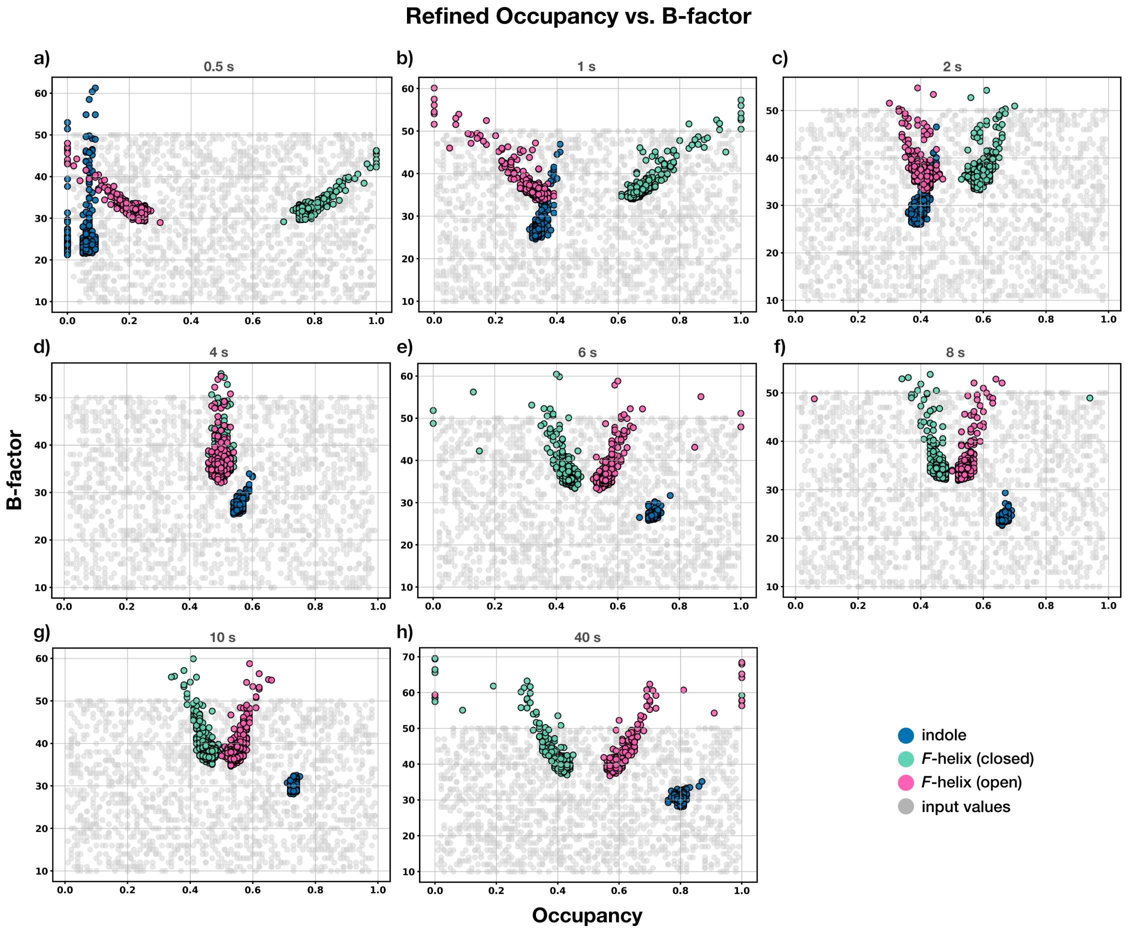


**Supplementary Figure 1 Occupancy–B-factor distributions for the full time series.**

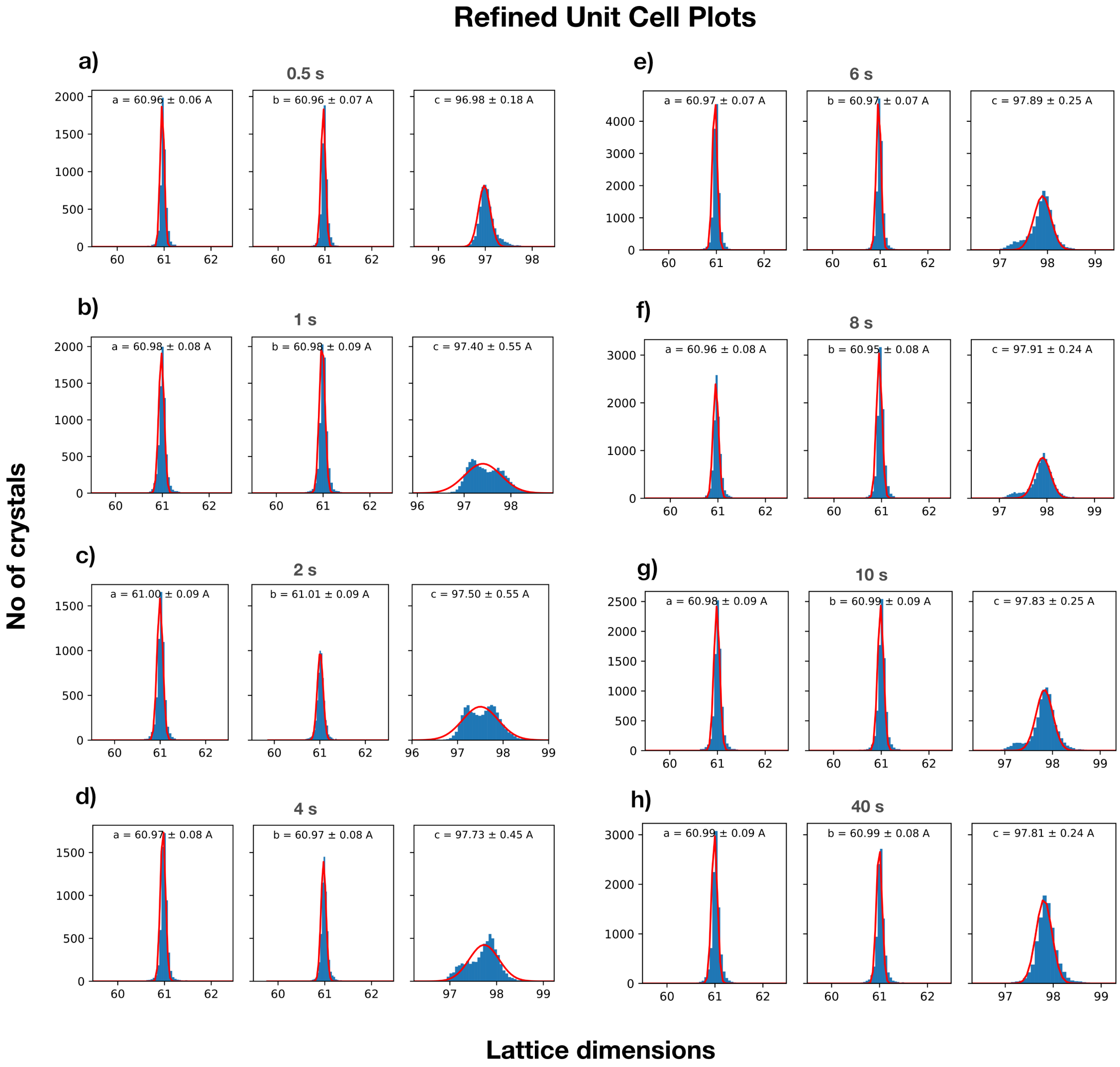


**Supplementary Figure 2 Refined unit cell parameter plots.** CrystFEL plots of the refined unit cell parameters for T4L-L99A crystals as a function of delay time. Blue bars correspond to the number of crystals refined for each value and the red curve to the gaussian fit, resulting in a mean value for each unit cell parameter.

### Data collection and refinement statistics

#### Supplementary Table 1

**Data collection and refinement statistics of T4L-L99A data.** *Values in the highest resolution shell are shown in parentheses.*

| Sample | apo | indole, 0.5 s | indole, 1 s | indole, 2 s | indole, 4 s | indole, 6 s | indole, 8 s | indole, 10 s | indole, 40 s | indole, long |
|---|---|---|---|---|---|---|---|---|---|---|
| PDB ID | 29NF | 29NG | 29NH | 29NI | 29NJ | 29NK | 29NL | 29NM | 29NN | 29NO |
| ***Data collection*** | | | | | | | | | | |
| Source | | | | | P14.2 (T-REXX) | | | | | |
| Beam size (μm) | | | | | 7 x 10 | | | | | |
| Flux (ph/s) | 1.0 x $10^{12}$ | 1.2 x $10^{12}$ | 1.2 x $10^{12}$ | 1.2 x $10^{12}$ | 1.0 x $10^{12}$ | 1.0 x $10^{12}$ | 1.2 x $10^{12}$ | 1.0 x $10^{12}$ | 1.0 x $10^{12}$ | 1.0 x $10^{12}$ |
| Energy (keV) | | | | | 12.7 | | | | | |
| Exposure time/ position (ms) | | | | | 5 | | | | | |
| Number of crystals | 16784 | 5403 | 6973 | 5511 | 5834 | 12618 | 8727 | 8229 | 9285 | 13855 |
| Space group | | | | | $P3_221$ | | | | | |
| Unit cell parameters/ (refined) | | | | | | | | | | |
| $a$, $b$, $c$ (Å) | 60.96, 60.98, 69.99 | 60.96, 60.96, 96.98 | 60.98, 60.98, 97.40 | 61.00, 61.01, 97.50 | 60.97, 60.97, 97.73 | 60.97, 60.97, 97.89 | 60.96, 60.95, 97.91 | 60.98, 60.99, 97.83 | 60.99, 60.99, 97.81 | 60.95, 60.98, 98.18 |
| $\alpha$, $\beta$, $\gamma$ (°) | | | | | 90, 90, 120 | | | | | |
| Resolution range (Å) | 52.74-1.70 (1.76-1.70) | 52.74-1.91 (2.03-1.91) | 52.74-1.91 (2.03-1.91) | 52.74-1.91 (2.03-1.91) | 52.74-1.91 (2.03-1.91) | 52.74-1.91 (2.03-1.91) | 52.74-1.91 (2.03-1.91) | 52.74-1.91 (2.03-1.91) | 52.74-1.91 (2.03-1.91) | 52.74-1.70 (1.76-1.70) |
| Total reflections | 5752833 (342851) | 1640703 (128263) | 2686852 (210839) | 2217676 (174403) | 2438035 (190991) | 5042938 (395364) | 2853767 (223055) | 2645989 (206529) | 2556637 (199717) | 4516792 (264407) |
| Unique reflections | 22682 (2231) | 23585 (2323) | 23585 (2325) | 23585 (2325) | 23585 (2323) | 23585 (2324) | 23585 (2323) | 23585 (2323) | 23585 (2325) | 23525 (2299) |
| Mean I/$\sigma$(I) | 6.61 (2.28) | 2.89 (1.59) | 3.25 (1.28) | 2.99 (1.21) | 3.28 (1.39) | 4.70 (2.15) | 3.56 (1.69) | 3.13 (1.02) | 2.82 (0.79) | 5.78 (1.87) |
| Completeness | 100 (100) | 100 (100) | 100 (100) | 100 (100) | 100 (100) | 100 (100) | 100 (100) | 100 (100) | 100 (100) | 100 (100) |
| Redundancy | 253.6 (153.7) | 69.6 (55.2) | 113.9 (90.7) | 94.0 (75.0) | 103.4 (82.2) | 213.8 (170.1) | 121.0 (96.0) | 112.2 (88.9) | 108.4 (85.9) | 192.0 (115.0) |
| $R_{split}$ | 0.141 (0.447) | 0.289 (0.652) | 0.220 (0.828) | 0.237 (0.882) | 0.209 (0.752) | 0.149 (0.479) | 0.206 (0.608) | 0.208 (1.015) | 0.226 (1.285) | 0.154 (0.533) |
| $CC^*$ | 0.993 (0.928) | 0.976 (0.851) | 0.989 (0.808) | 0.987 (0.785) | 0.990 (0.831) | 0.994 (0.925) | 0.989 (0.883) | 0.992 (0.746) | 0.991 (0.677) | 0.992 (0.904) |
| $CC_{1/2}$ | 0.971 (0.756) | 0.909 (0.567) | 0.958 (0.485) | 0.949 (0.445) | 0.962 (0.529) | 0.977 (0.748) | 0.958 (0.639) | 0.969 (0.386) | 0.966 (0.297) | 0.967 (0.691) |
| ***Refinement*** | | | | | | | | | | |
| Number of reflections/ (unique) | 22643 (2795) | 16691 (2715) | 16691 (2715) | 16682 (2709) | 16692 (2713) | 16691 (2713) | 16688 (2712) | 16682 (2712) | 16684 (2708) | 23488 (2868) |
| $R_{work}$ | 0.18 | 0.20 | 0.18 | 0.19 | 0.19 | 0.17 | 0.18 | 0.18 | 0.20 | 0.17 |
| $R_{free}$ | 0.20 | 0.23 | 0.20 | 0.21 | 0.21 | 0.20 | 0.21 | 0.20 | 0.22 | 0.19 |
| Occupancy - Indole | - | 0.07 | 0.34 | 0.39 | 0.54 | 0.74 | 0.65 | 0.75 | 0.81 | - |
| Occupancy - $F$-helix/ (closed/open) | - | 0.66/0.34 | 0.64/0.36 | 0.57/0.43 | 0.51/0.49 | 0.49/0.51 | 0.49/0.51 | 0.48/0.52 | 0.47/0.53 | - |
| Number of non-/ hydrogen atoms | 1395 | 1454 | 1464 | 1448 | 1442 | 1467 | 1460 | 1448 | 1443 | 1401 |
| Protein | 1312 | 1370 | 1370 | 1370 | 1370 | 1370 | 1370 | 1370 | 1375 | 1312 |
| Solvent | 83 | 75 | 85 | 69 | 63 | 88 | 81 | 68 | 58 | 79 |
| Ligand | - | 9 | 9 | 9 | 9 | 9 | 9 | 10 | 10 | 10 |
| Wilson B-factor (Å$^2$) | 20.65 | 23.23 | 26.64 | 26.40 | 26.30 | 25.74 | 24.63 | 29.04 | 31.70 | 21.03 |
| Mean B-factor (Å$^2$) | | | | | | | | | | |
| Overall | 24.20 | 24.33 | 29.20 | 27.71 | 26.90 | 28.37 | 25.86 | 33.30 | 34.59 | 24.11 |
| Protein | 23.71 | 24.13 | 28.89 | 27.54 | 26.75 | 27.83 | 25.52 | 32.97 | 34.50 | 23.65 |
| Solvent | 32.02 | 28.56 | 34.86 | 32.05 | 30.89 | 37.32 | 32.51 | 40.68 | 37.39 | 31.99 |
| Ligand | - | 18.60 | 22.64 | 20.53 | 20.80 | 22.30 | 18.70 | 28.83 | 30.88 | 22.01 |
| ***Model quality*** | | | | | | | | | | |
| *RMS deviations* | | | | | | | | | | |
| Bond length (Å) | 0.015 | 0.003 | 0.006 | 0.003 | 0.004 | 0.003 | 0.003 | 0.013 | 0.003 | 0.015 |
| Bond angles (°) | 1.61 | 0.55 | 0.76 | 0.52 | 0.65 | 0.56 | 0.54 | 1.20 | 0.53 | 1.50 |
